# Molecular adaptation in Rubisco: discriminating between convergent evolution and positive selection using mechanistic and classical codon models

**DOI:** 10.1101/073684

**Authors:** Sahar Parto, Nicolas Lartillot

## Abstract

Rubisco (Ribulose-1, 5-biphosphate carboxylase/oxygenase) is the most important enzyme on earth, catalyzing the first step of CO2 fixation in photosynthesis. Its molecular adaptation to C4 photosynthetic pathway has attracted a lot of attention. C4 plants, which comprise less than 5% of land plants, have evolved more efficient photosynthesis compared to C3 plants. Interestingly, a large number of independent transitions from C3 to C4 phenotype have occurred. Each time, the Rubisco enzyme has been subject to similar changes in selective pressure, thus providing an excellent model for convergent evolution at the molecular level. Molecular adaptation is often identified with positive selection and is typically characterized by an elevated ratio of non-synonymous over synonymous substitution rates (dN/dS). However, convergent adaptation is expected to leave a different molecular signature, taking the form of repeated transitions toward identical or similar amino acids.

Here, we use a previously introduced codon-based differential selection model to detect and quantify consistent patterns of convergent adaptation in Rubisco in Amaranthaceae. We further contrast the results thus obtained with those obtained under classical codon models based on the estimation of dN/dS. We find that the two classes of models tend to select distinct, although overlapping, sets of positions. This discrepancy in the results illustrates the conceptual difference between these models, while emphasizing the need to better discriminate between qualitatively different selective regimes, by using a broader class of codon models than those currently considered in molecular evolutionary studies.

## Introduction

Rubisco (Ribulose-1, 5-biphosphate carboxylase/oxygenase) is an enzyme that catalyzes the major step in carbon fixation in all photosynthetic organisms. It is the most abundant protein on earth [1], as it encompasses about 50% of soluble proteins in C3 plants [2]. During carbon fixation, Rubisco reacts with both CO2 and O2 as its substrate, with poor distinguishing ability. The carboxylase activity results in the incorporation of inorganic carbon into the metabolic C3 pathway, whereas the oxygenase activity boosts the photorespiration pathway. The latter prompts both energy consumption and CO2 loss. The evolution of C3 pathway goes back to 3 billion years ago when the atmosphere comprised high CO2 and low O2. In those conditions, photorespiration would rarely happen. However, under present atmospheric conditions (lower CO2 and higher O2 concentration), photorespiration can represent a significant proportion of the enzymatic activity of Rubisco, such that the efficiency of photosynthesis can be dropped by 40% under unfavorable climates like hot and dry conditions [3]. As a result, some plants have developed an evolved improvement to C3 pathway called C4 photosynthesis as an adaptation to these changes in the environment.

About 85% of plants use C3 photosynthetic pathway [3], whereas only about 5% are C4 plants [4]. The rate of photosynthesis is different in these groups, being much more efficient in C4 plants than in C3 species. C4 photosynthesis mostly evolved as an adaptation to intense light, high temperature and aridity [5]. Hence C4 plants dominate the grassland plants in harsh climates such as tropical, subtropical and warm regions [6]. C4 eudicots appeared on Earth much later than the first phototrophs.

The evolution of C4 plants from C3 ancestors consists of both anatomical and biochemical changes. These modifications allow C4 plants to concentrate more CO2 around Rubisco, such that the oxygenase activity and the subsequent photorespiration are partially or completely repressed. The kinetics of Rubisco has been altered in C4 plants, leading to lower specificity [7] and higher efficiency [8].

Interestingly, a relatively large number of independent transitions from C3 to C4 phenotype have occurred across monocots and eudicots. Each time, the Rubisco enzyme has been subject to similar selective pressure for tuning the tradeoff between substrate specificity and yield. As a consequence, C4 photosynthesis is an excellent model for convergent evolution at the molecular level in response to environmental changes [9]. In terms of applications, finding C4 plant features and applying them into C3 plants such as rice can be potentially used to increase crop yields [10, 11]. Considering the above issues, understanding how selection acts on Rubisco in C4 plants compared to C3 ancestors can be very beneficial.

Based on these considerations, the evolution of Rubisco has been paid a lot of attention in recent years [12-18]. Kapralov et al [14] tried to identify positive selection in Rubisco based on molecular phylogenetic analyses. Using codon models allowing for varying selection among sites (implemented in codeML [19]), they detected sites under positive selection in some photosynthetic organisms, especially in the main lineages of land plants. More recently, Kapralov et al [15] used a similar method to investigate the evolution of Rubisco in C4 plants in a large group of C4 eudicots and found sites under positive selection in C4 lineages. They observed that some of those positively selected sites (281 and 309) appear to display consistent patterns of amino acid substitutions associated with the C3 to C4 transition.

These empirical analyses raise an interesting question, concerning the use of codon models to characterize selective regimes in protein-coding sequences. Typically, elevated dN/dS, results from ongoing adaptive processes, by which a protein-coding gene is constantly challenged by ever-changing selective forces. However, in its general form, this process of ongoing adaptation needs not be associated with repeated transitions toward the same amino acid at a given position, independently across multiple lineages, and could instead constantly elicit new amino acids at positively selected sites. In contrast, the multiple transitions between the C3 and C4 photosynthetic regimes represent a case of *convergent* evolution. At the molecular level, this is expected to result in recurrent *directional* selection, thus, potentially favoring the same amino acid(s) at the same sites upon each C3-C4 transition. In addition, the overall dN/dS induced by this process of recurrent directional selection is fundamentally determined by the rate of C3/C4 transitions across the phylogeny, which may not be sufficiently high to induce a dN/dS greater than 1 at those positions that are susceptible to respond to this convergent evolutionary process. Thus, positive selection, like what is formalized by classical codon models (i.e. by an elevated dN/dS), may not be the most adequate selective regime to test in the present case.

Convergent amino acid substitutions which potentially linked to adaptation to the C4 phenotype have been more directly been investigated by Studer et al [18]. These authors used the TDG09 model, allowing for site- and condition-specific amino acid preferences [20], to identify sites under condition-dependent selection. Ancestral reconstructions were also performed and interpreted using a structural model of the protein, suggesting the existence of a conflict between the adaptive tuning of the enzymatic activity of the protein and the preservation of the protein stability.

The TDG09 model represents an approach for identifying consistent patterns of molecular adaptation caused by convergent evolution, thus representing a more adequate formalization of the selective regime induced by repeated transitions between C3 and C4 regimes. Recently, we have developed an approach similar to the TDG09 model, with the additional feature that our model is formulated at the codon level and uses a mechanistic derivation of the codon substitution process, using the mutation-selection formalism. The TDG09 model, in contrast, is phenomenological and is formulated directly at the amino acid level. One main advantage of our differential selection (DS) codon-based model is to offer a principled approach to tease out the respective contributions of mutation, selection pressure and random drift in the observed sequence patterns across plant groups. In addition, it is more directly comparable with codon-based analyses previously conducted on the evolution of Rubisco in monocots and eudicots.

Here, we re-assess the question of positive versus convergent selective patterns in the Rubisco gene in eudicots, using two types of codon models: first, we apply our DS model to identify amino acids which are differentially selected at specific positions along the Rubisco sequence, as a function of the photosynthesis pathway. Second, we implemented Bayesian versions of the classical dN/dS-based codon models, allowing for both site- and condition-specific modulations of the dN/dS ratio, and applied them to the Rubisco dataset. We find that the two classes of models tend to select distinct, although overlapping, sets of positions. Altogether, our analysis emphasizes the existence of qualitatively different adaptive regimes undergone by protein-coding genes, and the need to better discriminate between these distinct regimes by using a broader class of codon models than those currently considered in molecular evolutionary studies.

## Materials and Methods

### Sequence data, tree, and partitioning scheme

We obtained the Amaranthaceae rbcL multiple sequence alignment from Kapralov et al [15]. The dataset consists of 179 rbcL sequences of length 1341 base pairs (the first 22 coding positions are missing). Out of 179 sequences, 84 and 95 sequences belong to C4 and C3 species, respectively. The phylogenetic tree (Figure 1) was reconstructed using maximum likelihood inference, as in [15].

**Figure 1.**
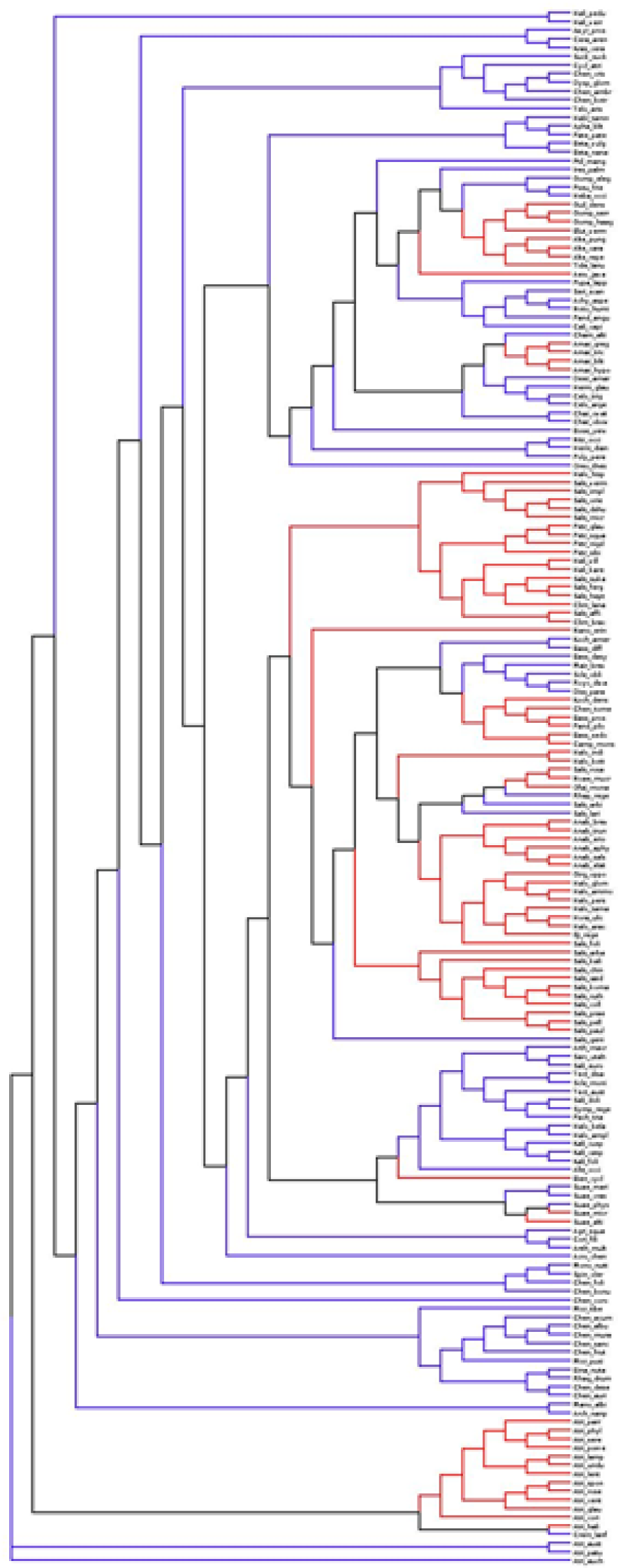
Amaranthaceae phylogenetic tree. The tree partitioned according to model M3 in C3 condition (blue), C4 condition (red) and interior branches (black).

The phylogenetic tree was partitioned according to two alternative schemes, with *K=3* or *K=2* distinct conditions, based on the type of the photosynthetic pathway. In the three-condition scheme, the largest monophyletic clades exclusively composed of C3 or C4 species were first identified and were defined as conditions 1 and 2. The branches at the base of each C3 and C4 clades were also included in conditions 1 and 2, respectively. All other branches outside from these clades were considered as belonging to condition 0. The model that employs this approach is called DS3 and its phylogenetic tree is illustrated in figure 1. The two-condition setup (model DS2) differs from the three-condition scheme (model DS3) by allocating all branches outside of the C4 monophyletic clades (together with their basal branches) by default to the C3 condition. Model DS2 amounts to assuming a maximum-parsimony reconstruction of the evolution of the photosynthetic regime, under the assumption that evolutionary transitions are exclusively from C3 to C4, with no reversion back to C3 [21]. However, model DS2 statistically implies a comparison between two conditions that are unevenly represented along the phylogeny, both in terms of total number of branches (152 for C4 versus 203 for C3) and concerning the evolutionary depth (the DS3 condition is mostly represented by recent branches, while the DS2 condition encompasses both ancient and recent lineages). In this respect, the advantage of model DS3 is to balance the empirical signal between the two conditions of interest (C3 and C4, represented by 158 and 153 branches under the DS3 model), and to focus exclusively on recent branches of the phylogeny for both conditions.

### Differential selection model

The principles of the differential selection model were introduced previously [22] and we only recall the general structure.

We use mutation-selection formalism, as in Halpern and Bruno [23] or Rodrigue et al [24]. According to this formalism, the substitution rates between codons are derived from first principles of population genetics, in terms of mutation rates and selective effects, the latter being explicitly modelled and assumed to operate exclusively at the level of the amino acid sequence.

More specifically, consider a sequence of *N* coding positions (*3N* nucleotide positions). The number of conditions across the phylogenetic tree is denoted as *K* (*K=2* or *K=3*, depending on the partition scheme, see above). The mutation process is assumed to be time-reversible and homogeneous among sites and across lineages. It is thus entirely characterized by a general time-reversible 4x4 matrix *Q*. In contrast, the selective forces acting at the amino acid level are both condition- and position-specific. Accordingly, for each position *i=1..N* and each condition *k=1..K*, we introduce an array of 20 non-negative fitness factors 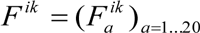, one for each amino acid. In the following, these 20-dimensional vectors will be referred to as amino acid *fitness profiles*. In the present version of the model, they are assumed to be random effects across sites and conditions, drawn *iid*. from a uniform Dirichlet distribution.

Once these mutation rates and fitness factors are specified, the substitution process can be defined as follows. Consider the substitution rate between codon *c*_1_ to *c*_2_ at site *i* and condition *k*, where codons *c*_1_ and *c*_2_ are assumed to vary only at one nucleotide position, with respective nucleotide states *n*_1_ and *n*_2_ at that position, and to encode for amino acids *a*_1_ and *a*_2_, respectively. First, we define a scaled selection coefficient, associated with codon *c*_2_, seen as a mutant in the context of a population in which the wild-type allele is *c*_1_. Since selection is assumed to act only at the level of the amino acid sequence, this scaled selection coefficient is given by

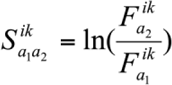

Then, the rate of substitution between codon *c*_1_ and *c*_2_ is given by the product of the mutation rate and the relative fixation probability *P* (i.e. relative to neutral). This fixation probability is itself dependent on the scaled selection coefficient just defined. Using the classical diffusion approximation, this relative fixation probability can be expressed as

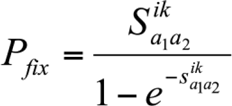

Thus, finally, the rate of substitution between codons *c*_1_ and *c*_2_ at position *i* and under condition *k* is given by

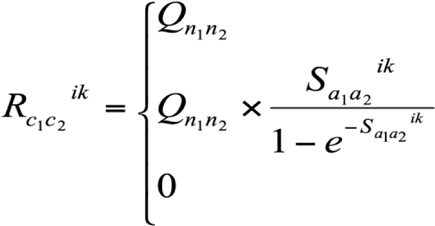

### Omega-based codon models

As an alternative to mutation-selection models, one of the most well-known and widely used methods for characterizing the selective regimes, involved in the evolution of protein-coding genes, is to estimate the ratio of non-synonymous (dN) and synonymous (dS) substitution rate, denoted as ω (=dN/dS). These omega-based models were first proposed by Goldman and Yang [25] and Muse and Gaut [26], and subsequently complexified to account for site- and branch-specific modulations of the dN/dS ratio [27-30].

Here, we use the Muse and Gaut formalism, and propose a Bayesian model allowing for site- and condition-specific modulations of ω=dN/dS. According to this model, the instantaneous substitution rate from codon *c*_1_ to *c*_2_ at site *i* and condition *k* is now specified as follows

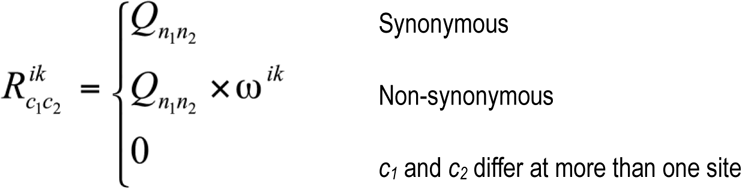

Here, *ω^ik^* is thus the dN/dS ratio for site *i* and under condition *k*. For each condition *k*, the *ω^ik^* s, for *i=1…N*, are modelled as random effects across sites, drawn *iid*. from a gamma distribution of shape and scale parameters *α^k^* and *β^k^*.

We consider two alternative versions of this omega-based model: in model OM1, we assume only one condition, thus defining a single (global) value of *ω^i^* across the whole phylogenetic tree for site *i*; in model OM3, on the other hand, the tree is partitioned into three conditions according to the photosynthesis pathways, exactly as for model DS3 above, and a distinct value *ω^ik^* is allowed for site *i* and under condition *k=1…3*.

**Priors-** In all analyses presented below, the topology (*τ*) of the tree is fixed. For all models, the prior on branch lengths is a product of independent Exponentials of mean *λ*; the hyperparameter *λ* is from an Exponential distribution of mean 0.1; the prior on relative exchangeabilities of the mutation process is a product of Exponentials of mean 1; on the mutational equilibrium frequency vector, it is a uniform Dirichlet distribution. As mentioned above, under the DS2 and DS3 models,the site- and condition-specific fitness profiles, 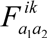, are random effects integrated over a Dirichlet distribution. Concerning the OM1 and OM3 models, the site-and condition-specific dN/dS values (*ω^ik^*) are random effects integrated over a gamma distribution of shape and scale parameters *α^k^* and *β^k^*. For each *k=1…K*, we assume an exponential prior of mean 1 for *α^k^* and *β^k^*.

### MCMC sampling

To sample the parameters from their joint posterior distribution, we used the general MCMC approach previously described in [22, 31, 32]. This approach consists of an alternation between stochastic mapping of the detailed substitution history at each coding site, followed by a long series of Metropolis-Hastings updates of all parameters and all random-effects across sites and across conditions, conditional on this stochastic mapping. After 400 points of burn-in have been removed, posterior estimates are obtained by averaging over the remaining of the MCMC chain.

### Post-analysis

Under the DS models, for a given configuration of the model (typically drawn from the posterior distribution by MCMC), differential selection between two conditions C3 and C4 is simply calculated as the log-ratio between the amino acid fitness profiles ascribed to conditions 1 and 2

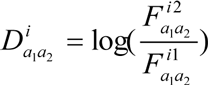

These arrays of 20 differential selection effects (for the 20 amino acids) at each position are then averaged over the posterior distribution by MCMC. A position is deemed to show strong statistical support for a differential effect in favor of amino acid *a*_2_ (in condition C4) over amino acid *a*_1_ (in condition C3) if the posterior probability that 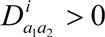 is greater than 0.90.Conversely, strong support for a negative differential effect (i.e. a differential effect against *a*_2_ in favor of *a*_1_) is called whenever the posterior probability that 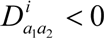 is greater than 0.90.

Under the OM models, the posterior mean value of site- and condition-specific dN/dS is reported. Position *i* is deemed to have a strong support for positive selection under condition *k* if the posterior probability that *ω^ik^* > 1 is greater than 0.90.

## Results

Amaranthaceae is one of the families with the largest number of C4 plant species. This makes it a suitable case for Differential Selection (DS) analysis. Based on a multiple sequence alignment of rbcL genes and an annotated phylogenetic tree of Amaranthaceae, our DS model captures site-specific amino acid preferences as vectors of 20 fitness factors (for the 20 amino acids) under each condition. Then, contrasting for each position, the fitness factors estimated in the two conditions of interest (here, in the C3 and C4 regimes), allows us to identify positions for which the fitness of a specific amino acid has undergone a significant change, either upward or downward, associated with the transition between the C3 and the C4 photosynthetic regime (see Methods).

### DS2 versus DS3: finding an optimal contrast between C3 and C4

As described in the methods, the tree was divided in either two or three conditions, resulting in two version of the differential-selection codon model, referred to as DS2 and DS3. In DS2, the interior branches (black branches in figure 1) which connect C3 and C4 clusters are defined as C3. This corresponds to a plausible reconstruction of ancestral photosynthetic regimes across the group, as no reversal from C4 to C3 is known [21].

The selection profiles at position 306-331 estimated by DS2 are shown in figure 2. In this figure, we use a graphical logo representation [33] to display both absolute and differential fitness distributions. Absolute logos for the reference condition represent the fitness of amino acids under a specific condition, with the height of the letter being proportional to the fitness of the corresponding amino acid. Differential logos, on the other hand, represent the difference in log-fitness between two conditions: letters above (resp. below) the line corresponds to amino acids whose fitness is increased (resp. decreased) in each condition, compared to its parent condition.

**Figure 2.**
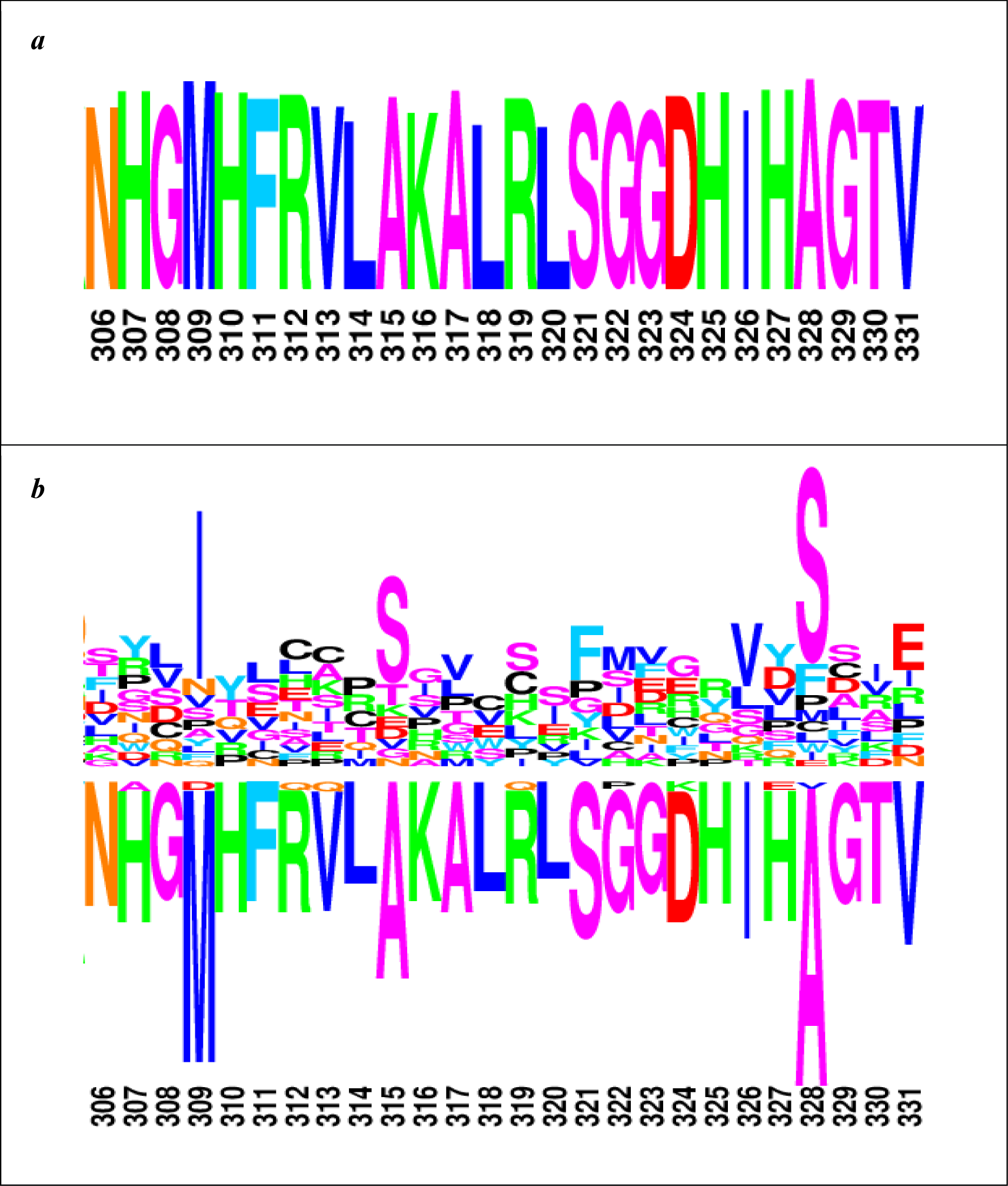
Global and differential selection profiles (differential for C4), for position 306-331, by model DS2.

The baseline selection profile (figure 2-a) captures the absolute amino acid fitness for C3 plants. This profile primarily reflects the strong conservation of the protein sequence, with one single amino acid overwhelmingly favored at most positions. The differential profile between C4 and C3 (figure 2-b) shows interesting patterns of opposite selective effects concerning pairs of amino acids, specifically at positions 309, 315 and 328. However, the differential profile between the C4 and C3 is also characterized by an inferred background of apparently non-specific differential selective effects concerning all major amino acids represented in the absolute fitness profile under C3: essentially, the absolute profile under C3 displays the consensus sequence of the alignment, while the differential profile between C4 and C3 reproduces this consensus sequence, although now below the line. This is likely to be a statistical artifact, which might have two alternative explanations. The first one is the possible existence of non-fixed polymorphic states in the multiple sequence alignment. These mutations, whose fate is to be ultimately removed by purifying selection, are expected to be mapped specifically along the terminal branches of the phylogeny, and may thus contribute to an apparent decrease in the inferred fitness of ancestral amino acids in the condition that is most enriched in terminal branches (here, C4). As another possible explanation, the number of branches allocated to the C4 condition is smaller than that allocated to the C3 condition, potentially leading to a difference in statistical power between the two conditions. As a result, and in the presence of shrinkage mediated by the prior, the fitness of conserved amino acids is inferred to be higher in that condition that is endowed with the largest number of branches (here, C3).

One way to avoid this artifact is to balance the signal between conditions C3 and C4, by allocating the interior branches of the tree to another baseline condition and by restricting the inference of C3-specific selection to the monophyletic groups of C3 species. In the present case, there are a comparable number of C3 and C4 branches (about 150 for each). This new setting (model DS3) is therefore expected to result in a much more balanced assessment of the differential selection effects between recent C3 and C4 lineages.

Indeed, and unlike the differential profile between C3 and C4 provided by the DS2 model (figure 2), the differential profile given by DS3 between recent C3 and C4 lineages (figure 3) appears to have more reasonable properties: sparse, selecting a small number of positions for which specific amino acids appear to be differentially selected between the two photosynthetic regimes, it is also balanced between positive and negative effects (above and below the line, respectively). We see for instance that, at position 309, the fitness of Methionine is substantially decreased in C4 plants, compared to C3 species (p.p. =0.93). Correlatively, the fitness of Isoleucine is increased at that position (p.p. = 0.87). Similarly, residue 328 is identified by the DS3 model as being the position of the rbcL gene under the highest differential selection effect between C3 and C4 Amaranthaceae species. At site 328 Alanine is globally preferred in Amaranthacea eudicots, yet in C4 group, it’s fitness is significantly decreased (p.p. =0.99) in favor of Serine, whose fitness is increased compared to what prevails in C3 lineages (p.p.=0.96). Based on these observations, in the following, we conduct all differential selection analyses under the DS3 model. The complete C4/C3 differential selection logo, for the whole sequence alignment, is displayed in the appendix.

**Figure 3.**
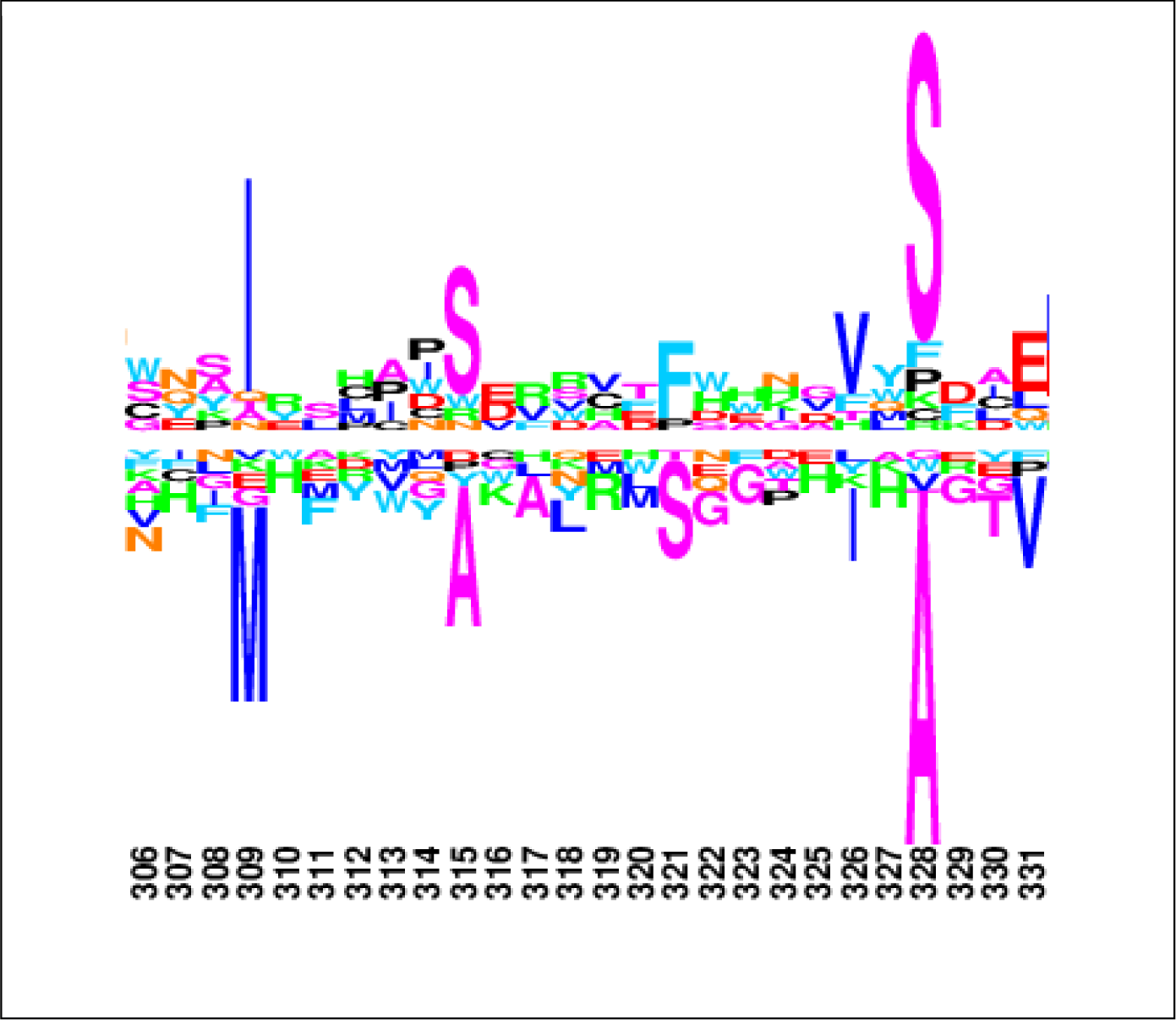
C4/C3 differential selection profile for position 309-328, under the DS3 model.

### Mutation-selection (DS) versus omega-based (OM) codon models

In addition to DS3, which belongs to the family of mutation-selection codon models (Halpern and Bruno style), the Amaranthacea dataset was also analyzed under two omega-based models (Muse and Gaut style, see Methods). The first of these models (OM1) is a site-specific model: each site has its own value for ω=dN/dS, all of which are modeled as site-specific gamma-distributed random-effects. By selecting sites with a high posterior probability of having a value of dN/dS greater than 1, model OM1 allows for the detection of sites under positive selection globally across Amaranthacea dataset. The second model (OM3) allows for independent values of the dN/dS simultaneously across sites and conditions. Conditions are defined as in the DS3 model (internal branches, as well as terminal C3 and C4 clades). Thus, OM3 unlike OM1 allows for the detection of sites under positive selection specifically in C3 or in C4 species. To facilitate the comparison, all three models, DS3, OM1 and OM3, were implemented in a Bayesian framework, using similar strategies for designing the models (both dN/dS and differential selective effects modelled as either global or condition-dependent *iid* random-effects across sites) and for detecting significant effects (based on the posterior probability for a site to have a value of ω> 1 or a differential selective effect greater and smaller than 0, globally or in a given condition). The results of these analyses are summarized in Table 1. In this table, all sites for which a strong support (p.p. >0.90) was found under at least one of the three models are reported.

**Table 1.**
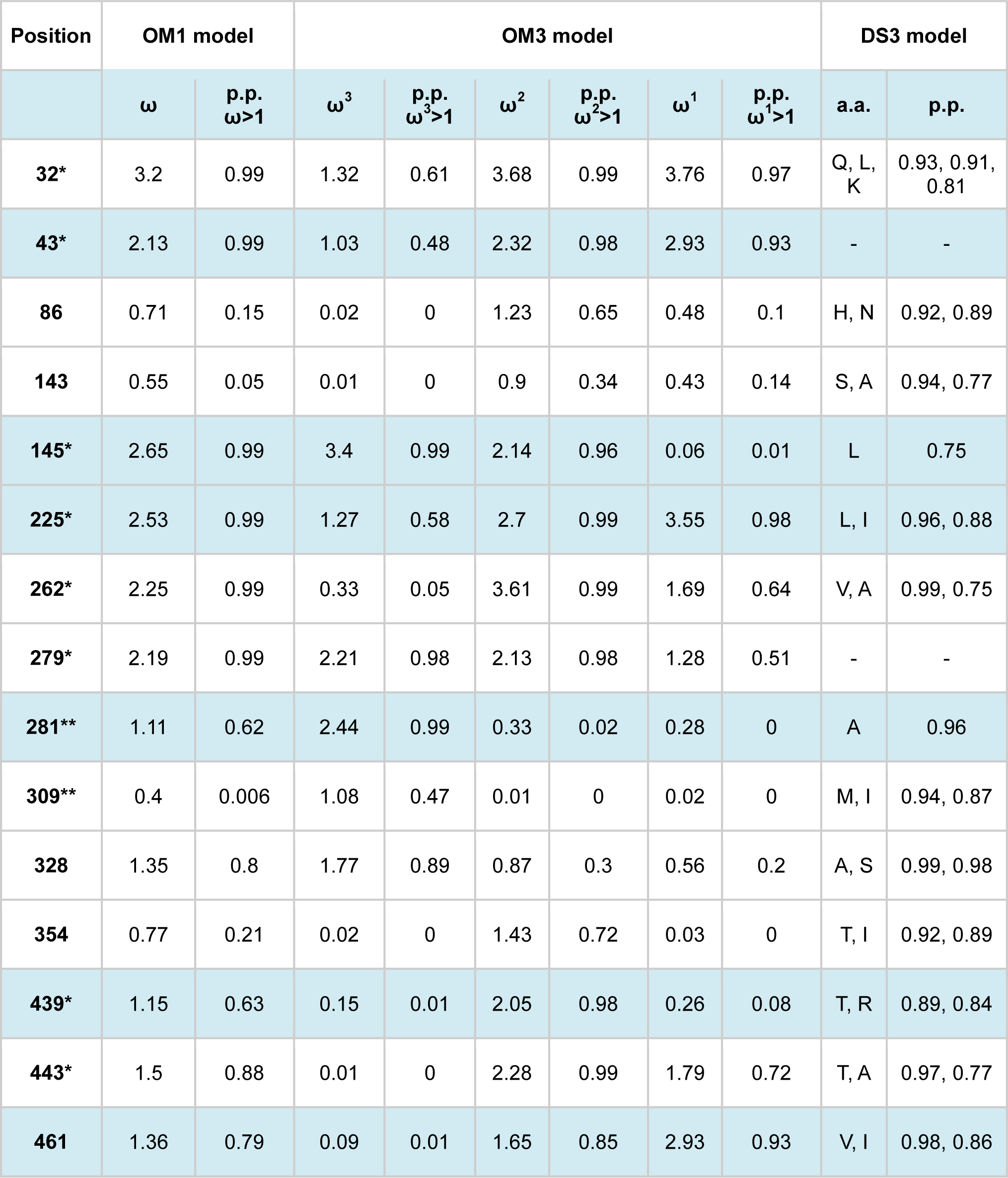
Findings of OM1, OM3 and DS3 model with their posterior probabilities. Positions with one and two asterix are those found previously by Kapralov et al (site and branch-site models, respectively).

| Position | OM1 model |  | OM3 model |  |  |  |  |  | DS3 model |  |
| --- | --- | --- | --- | --- | --- | --- | --- | --- | --- | --- |
| | $\omega$ | p.p.<br>$\omega > 1$ | $\omega^3$ | p.p.<br>$\omega^3 > 1$ | $\omega^2$ | p.p.<br>$\omega^2 > 1$ | $\omega^1$ | p.p.<br>$\omega^1 > 1$ | a.a. | p.p. |
| **32\*** | 3.2 | 0.99 | 1.32 | 0.61 | 3.68 | 0.99 | 3.76 | 0.97 | Q, L, K | 0.93, 0.91, 0.81 |
| **43\*** | 2.13 | 0.99 | 1.03 | 0.48 | 2.32 | 0.98 | 2.93 | 0.93 | - | - |
| **86** | 0.71 | 0.15 | 0.02 | 0 | 1.23 | 0.65 | 0.48 | 0.1 | H, N | 0.92, 0.89 |
| **143** | 0.55 | 0.05 | 0.01 | 0 | 0.9 | 0.34 | 0.43 | 0.14 | S, A | 0.94, 0.77 |
| **145\*** | 2.65 | 0.99 | 3.4 | 0.99 | 2.14 | 0.96 | 0.06 | 0.01 | L | 0.75 |
| **225\*** | 2.53 | 0.99 | 1.27 | 0.58 | 2.7 | 0.99 | 3.55 | 0.98 | L, I | 0.96, 0.88 |
| **262\*** | 2.25 | 0.99 | 0.33 | 0.05 | 3.61 | 0.99 | 1.69 | 0.64 | V, A | 0.99, 0.75 |
| **279\*** | 2.19 | 0.99 | 2.21 | 0.98 | 2.13 | 0.98 | 1.28 | 0.51 | - | - |
| **281\*\*** | 1.11 | 0.62 | 2.44 | 0.99 | 0.33 | 0.02 | 0.28 | 0 | A | 0.96 |
| **309\*\*** | 0.4 | 0.006 | 1.08 | 0.47 | 0.01 | 0 | 0.02 | 0 | M, I | 0.94, 0.87 |
| **328** | 1.35 | 0.8 | 1.77 | 0.89 | 0.87 | 0.3 | 0.56 | 0.2 | A, S | 0.99, 0.98 |
| **354** | 0.77 | 0.21 | 0.02 | 0 | 1.43 | 0.72 | 0.03 | 0 | T, I | 0.92, 0.89 |
| **439\*** | 1.15 | 0.63 | 0.15 | 0.01 | 2.05 | 0.98 | 0.26 | 0.08 | T, R | 0.89, 0.84 |
| **443\*** | 1.5 | 0.88 | 0.01 | 0 | 2.28 | 0.99 | 1.79 | 0.72 | T, A | 0.97, 0.77 |
| **461** | 1.36 | 0.79 | 0.09 | 0.01 | 1.65 | 0.85 | 2.93 | 0.93 | V, I | 0.98, 0.86 |

Under model OM1, 8 positions were found to have a dN/dS > 1 with a posterior probability greater than 0.90. These 8 positions are exactly those reported by Kapralov et al [15], found using the BEB approach implemented in CodeML. Note that the approach used here and the one implemented in CodeML are rather different in their statistical strategy for detecting sites under positive selection. The approach of CodeML relies on a mixture model, whereas the present approach explicitly assumes independent values of dN/dS across sites. The results obtained here therefore, suggest that at least in the present context, the details of the overall statistical strategy do not have a strong influence on the outcome of the model.

Kapralov et al [15] also reported an additional two sites, 281 and 309, detected by the branch-site model, and thus inferred to be under positive selection specifically in the C4 condition. Here, using the OM3 model, which allows for condition- and site-specific values for dN/dS, we found statistical support for positive selection in C4 only for position 281. For position 309, the posterior mean value of omega in condition C4 is indeed greater than 1 (1.04), although only with a weak posterior probability support (p.p. = 0.47). Conversely, it is worth noting that several sites (32, 43, 225, and 262) are inferred by model OM3 to be under positive selection only under C3, but not under C4.

Finally, the DS3 model uncovers a series of 11 sites under differential selection between C3 and C4 with p.p. > 0.90. These 11 sites include 4 of the sites discovered by model OM1, thus inferred to be under global positive selection (32, 225, 262 and 443), as well as sites 281 and 309 (inferred to be under positive selection specifically in C4, either by model OM3 or by branch-site models of CodeML). Conversely, and importantly, half of the discoveries made by the DS3 model (6 sites out of 11, including site 309) do not show any signal of positive selection under either OM1 or OM3.

## Discussion

### Differential selection patterns in Amaranthacea

Here we studied the molecular adaptations associated with the C3 to C4 transition in Amaranthaceae eudicots. Using a mechanistic codon model for detecting differential selection patterns associated with these adaptations, we found 11 positions to be under differential selection pressure between C3 and C4 eudicots. The amino acid substitutions undergone by these positions might have a conformational role in Rubisco enzyme in C4 plants, leading to its higher efficiency. Alternatively, they might be a compensatory mutation selected to maintain its optimized function.

For instance, residue 328, with the highest differential effect, locates in the active loop 6 of the enzyme. Replacement of hydrophobic A with polar S destabilizes the active site, which leads to more flexibility of its opening and closing [34] and might explain the higher efficiency of C4 plants. Site 281 lies in the core of C-terminal domain, and it may have a long-range effect on active loop 6 [18]. Position 309 is in the interface of C-terminal domains of two subunits within a dimer [18], which might have an effect on flexibility. Although residues 86, 354 and 461 are found to be under strong differential selection pressure between C3 and C4 Amaranthaceae, their exact role has not been yet specified. Position 461 locates near a large subunit residue (residue 466) which might account for the interaction with Rubisco activase.

### Comparing differential-selection and omega-based codon models

Previously, some positions have been found by other phylogenetic methods to be under specific selective regimes, potentially associated with the C3 to C4 transition. In particular, Kapralov et al [15] used the concept of dN/dS as selection strength along the coding sequence. Using classic dN/dS codon models, they uncovered a set of 10 positions putatively under positive selection, either globally over the tree (8 positions) or specifically in the C4 groups (2 positions). In order to further explore this point, we implemented new dN/dS codon models, allowing for site- and condition-specific dN/dS, in our Bayesian framework. Selecting sites based on the posterior probability support for dN/dS>1, we essentially recover the same set of positions as that reported by Kapralov et al (except for one position). On the other hand, if we compare the set of findings under dN/dS models and the differential mutation-selection model, we see only a partial overlap. Specifically, only half of the positions inferred to be under differential selection between C3 and C4 were also found by dN/dS models. Conversely, 4 of the 10 findings under both classes of dN/dS models show differential selection effects.

The partial overlap between the findings of omega-based and differential-selection models illustrates the conceptual difference between these models, and the fact that they are meant to capture fundamentally different selective patterns. Classic omega-based codon models are meant to detect an overall *acceleration* of the rate of non-synonymous substitution. Such accelerations are typically caused by what could be called *ongoing adaptation*, due to diversifying selection, ecological red-queens, or fluctuating selection caused by environmental changes. In contrast, differential-selection models are meant to capture convergent patterns of *directional selection* associated with a specific change in the environment, having occurred several times independently across the phylogeny.

These two classes of selective patterns are not completely mutually exclusive. In principle, recurrent substitution events due to directional selection caused by repeated transitions from C3 to C4 photosynthesis across the Amaranthacea family could result in an overall increase in the dN/dS observed at the corresponding sites. However, if the rate of C3 to C4 transitions is not sufficiently high, the resulting increase in dN/dS may not be enough to lead to a situation where dN/dS>1. As a result, some of the important condition-specific adaptations might be missed by dN/dS codon models. For instance, as illustrated here, positions 86, 143 or 354, which show a strong differential selection effect, yet have a dN/dS not exceeding 1.

In addition, this phenomenon of recurrent directional selection linked to repeated C3 to C4 transitions cannot explain that most of the sites inferred to be under positive selection have a dN/dS>1 globally over the tree, and often (e.g. positions 32, 43 and 279) even specifically within the C3 terminal clades, in which no such substitution event induced by C3 to C4 transition is supposed to have occurred. Concerning positions 43 and 279, for instance, no differential selection effect is detected by the DS3 model, while the dN/dS is inferred to be of the order of 2, including within terminal C3 clades. Thus the most likely explanation for the pattern of Darwinian evolution at those sites is simply the presence of ongoing adaptation that would not be directly related to the repeated transitions between C3 and C4 photosynthetic regimes.

Conversely, we observed some positions (in particular 262 and 461) that show a differential selection effect between C3 and C4, combined with a pattern of positive selection over the tree, *except* in the C4 condition, in which the dN/dS is specifically and markedly decreased (posterior mean dN/dS = 0.33 and 0.09, respectively). A possible explanation for this pattern would be that, in C3 species, those positions are available for ongoing adaptation to a constantly fluctuating environment, but that the transition to C4 photosynthesis essentially locks those positions into more specific adaptive amino acid states, thereby stopping the flux of adaptive substitutions at those sites. Of note, however, this conjunction of positive selection and differential selection effects (e.g. positions 32, 225) is not so easily explained in the context of the mutation-selection modeling framework used here. Mutation-selection models predict that the dN/dS should always be less than 1 at mutation-selection balance [35]. In the present case, this means that the DS model does not predict dN/dS greater than 1, except possibly during the transient phases following a change between the C3 and the C4 regimes -- thus, at any rate, not within the C3 clades.

Finally, Studer et al [18], analyzing a dataset of monocot families [12], reported a set of 18 differentially selected positions in monocots, using TDG09 model [20]. The approach behind the TDG09 model is similar to the one used here, with the main differences being that the DS model is formulated at the codon level (whereas the TDG09 model is expressed at the amino acid level) and the use of a Bayesian, rather than a maximum likelihood, statistical paradigm. Overall, there are 8 positions that are in common between their and the present analysis (328, 262, 281, 225, 143, 309, 251 and 228). These common positions can be declared as sites which are under condition-specific selection in monocots and eudicots. To have a better comparison of the two alternative approaches, however, it can be suggested to apply DS model on their monocot dataset presented by Christin et al [12].

## Conclusion

Rubisco has long been known to be under positive selection. In addition, it has been shown to evolve differently, and according to consistent molecular patterns, in C3 and C4 plants. The complex molecular evolutionary patterns displayed by the Rubisco gene in eudicots therefore represent an interesting case-study for assessing and comparing current codon modeling strategies. In this respect, our comparative analysis, by making an inventory of the amino acid-positions in rbcL sequences that are positively or differentially selected in C3 and C4 Amaranthaceae family, emphasizes the fundamental difference, in scope and meaning, between the two main classes of models currently considered in the literature: on one side, classic codon models based on the measure of the overall dN/dS, whose focus is primarily on positive selection; and on the other side, differential selection models, whose aim is instead, to detect convergent patterns of directional selection associated with repeated transitions between known evolutionary regimes. Our analysis also emphasizes that none of the models considered here, either omega-based models or mutation-selection approaches, offers a completely satisfactory explanation of the complex patterns of molecular evolution observed in Amaranthacea, and probably also present in other species groups – thus suggesting that further developments are still needed on the front of phylogenetic codon-models.

## Appendix

The first 21 amino acids are missing.

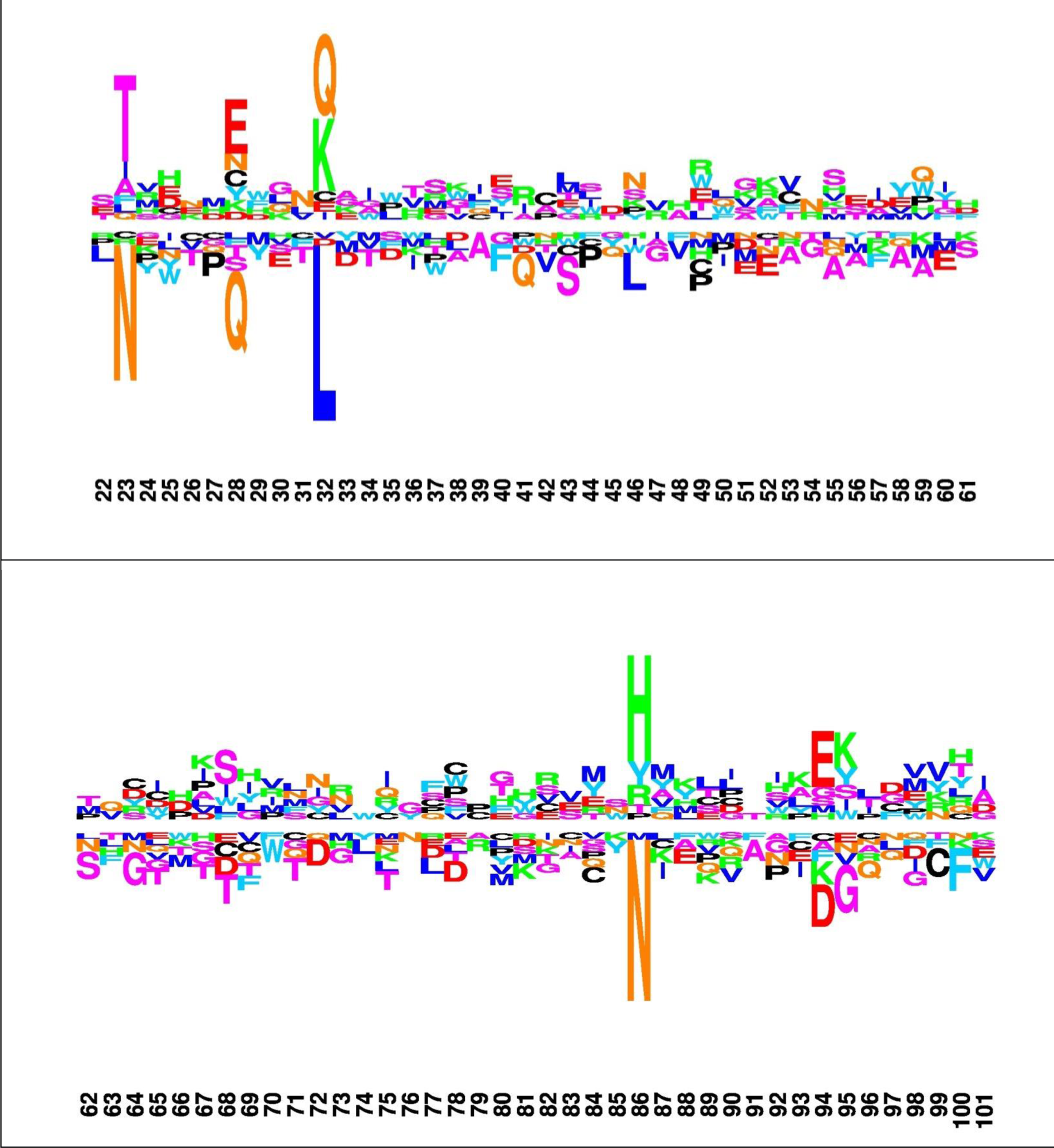

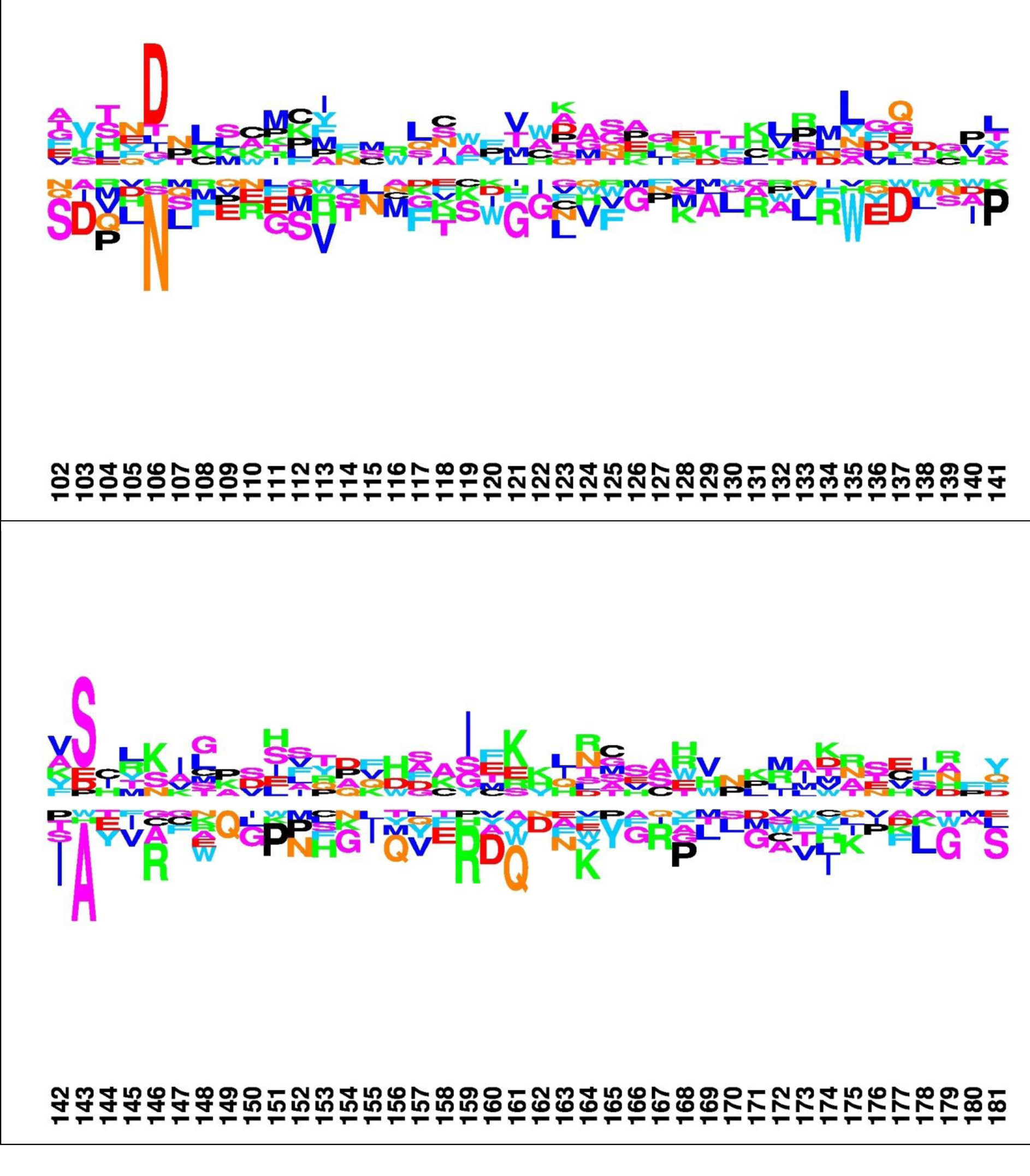

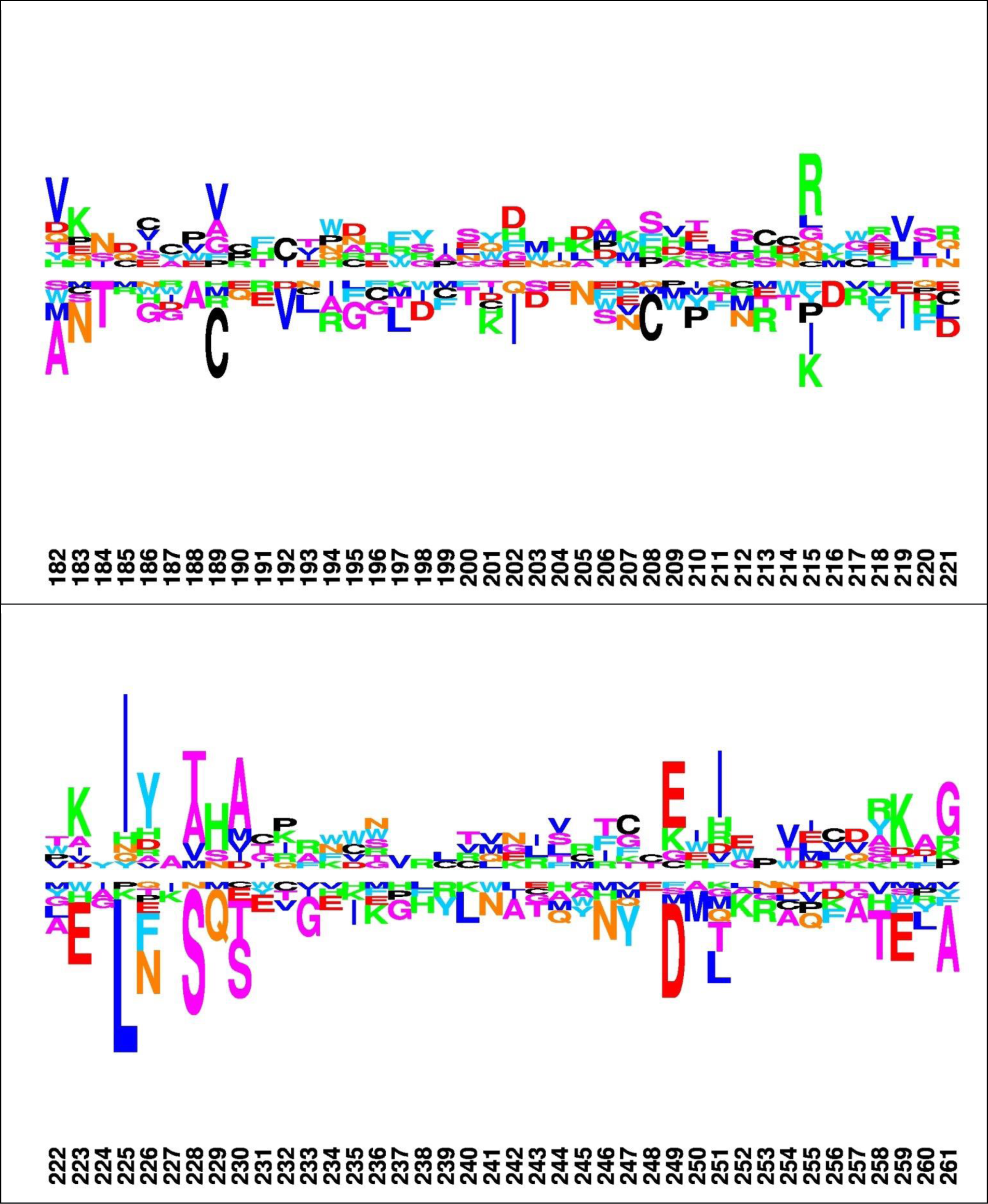

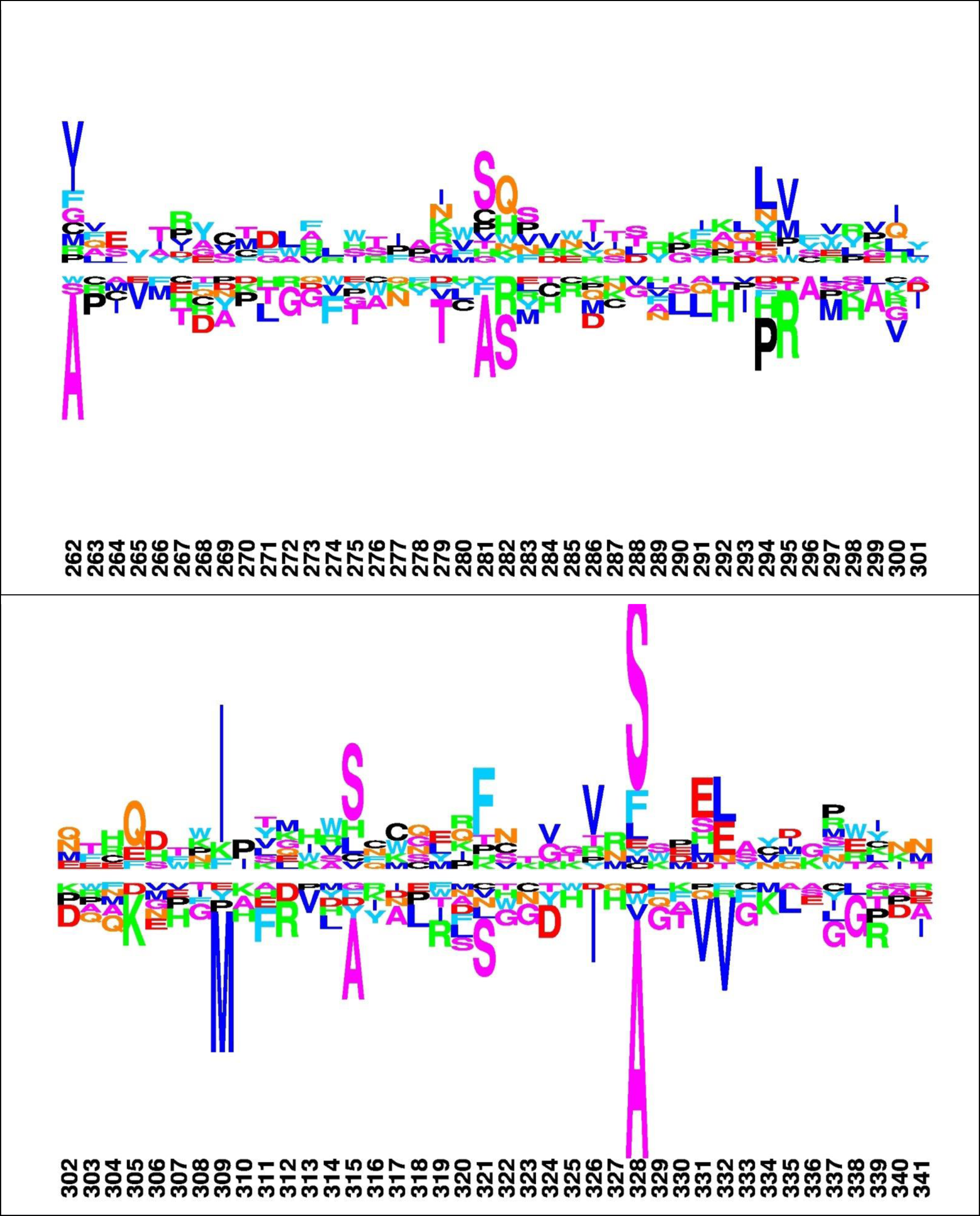

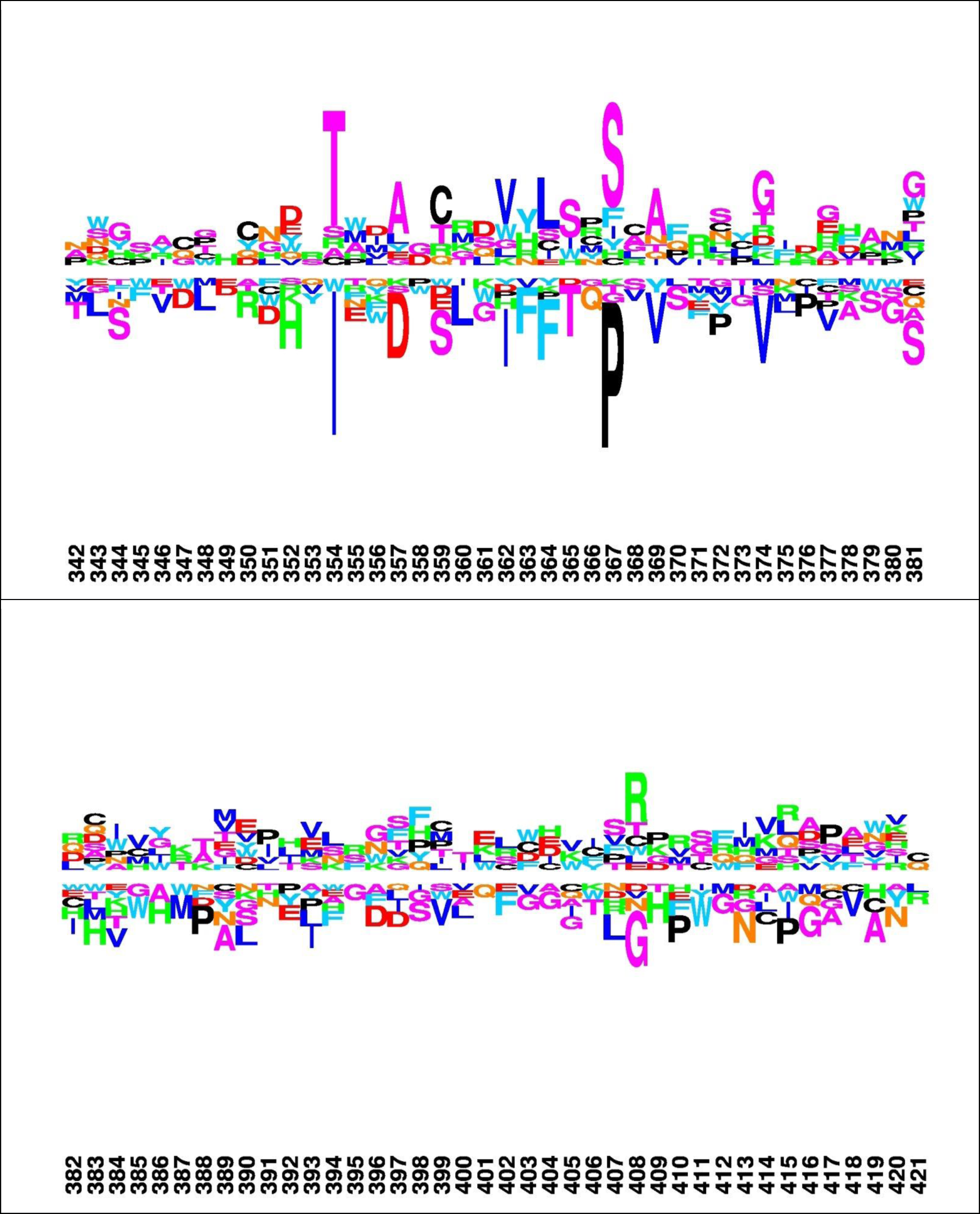

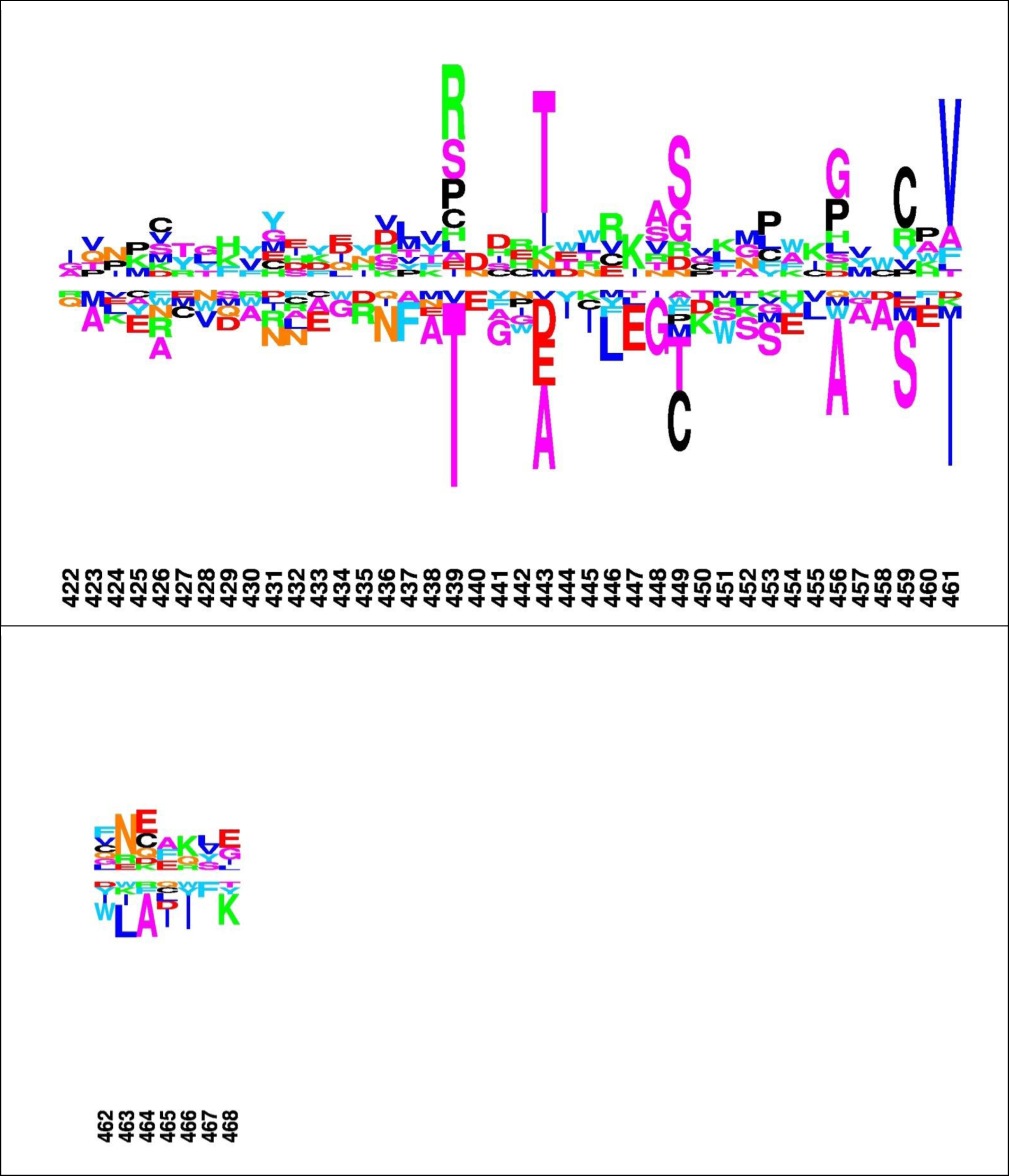

